# Characterization of a major thrashabilly locus in tetraploid wheat

**DOI:** 10.64898/2026.03.30.715257

**Authors:** Yael Lev-Mirom, Raz Avni, Moran Nave, Sharon Kulikovsky, Leah Oren, Tamar eilam, Hanan Sela, Assaf Distelfeld

**Affiliations:** Department of Evolutionary and Environmental Biology, Faculty of Natural Sciences and the Institute of Evolution, University of Haifa, Haifa, Israel

**Keywords:** wheat domestication, glume tenacity, threshability, QTL mapping, near-isogenic line, wild emmer wheat, *Tg1-B*, tetraploid wheat

## Abstract

The transition from hulled to free-threshing grain was a pivotal event in wheat domestication, enabling efficient harvesting and processing. Threshability in tetraploid wheat is controlled primarily by the *Q* locus and two *Tenacious glume* (*Tg*) loci on chromosomes 2A and 2B, yet the molecular basis of the major *Tg1-B* locus remains incompletely characterized. Here, we phenotyped a durum wheat x wild emmer wheat (WEW) recombinant inbred line (RIL) population across two field environments and performed QTL analysis for glume tenacity (TG), threshability ratio (THRR), and seed number per spike (SDNPS). A total of 19 significant QTLs were detected across six chromosomes. The largest-effect loci for both TG and THRR co-localized on chromosome 2B, with LOD scores up to 14.22 and phenotypic variance explained up to 31.2%, corresponding to the previously described *Tg1-B* locus. To validate this QTL, the donor RIL was backcrossed three times to Svevo to generate a near-isogenic line, NIL-65 (BC_3_F_5_), confirmed by whole-genome skim sequencing to carry a homozygous WEW introgression at *Tg1-B*. A segregating BC_4_F_2_ population derived from NIL-65 confirmed that plants homozygous for the dominant *Tg1-B* allele displayed significantly higher glume tenacity and intact glume morphology compared to *tg1-B* sister lines, which exhibited basal glume cracking characteristic of the free-threshing phenotype. Genotyping-by-sequencing delimited the causal interval to an approximately 11 Mb introgression on chromosome 2B. These results confirm the major role of *Tg1-B* in determining glume tenacity in tetraploid wheat, provide a validated near-isogenic germplasm resource, and lay the foundation for fine-mapping and functional characterization of the underlying gene(s).

## 1. Introduction

Wheat (genus: *Triticum*) is one of the most important crop species, essential for human nutrition from the Neolithic era until today, providing approximately 20% of global caloric intake. Wild emmer wheat (WEW; *Triticum turgidum* subspecies (ssp.) *dicoccoides* (Körn) Thell (2n=4x=28), AABB genomes), the progenitor of all wheat species, evolved about 0.5 million years ago from hybridization between the diploid *Triticum urartu* (genome AA) and a diploid *Aegilops* species (genome BB). The domesticated emmer wheat (DEW; *T. turgidum* ssp. *dicoccum*, genome BBAA) subsequently evolved to durum wheat (*T. turgidum* ssp. *durum*) and other forms of cultivated tetraploid wheat. Hexaploid bread wheat (*T. aestivum*, genome BBAADD) arose from the hybridization of domesticated tetraploid wheat with the diploid *Aegilops tauschii* (genome DD) (Feldman & Levy, 2023) Since Aaronsohn’s 1910 discovery of wild emmer wheat (WEW) in Rosh-Pina, it has been recognized as a vital resource for wheat improvement, offering extensive natural genetic variation and, together with domesticated durum wheat, serving as an increasingly tractable platform for dissecting domestication genetics and advancing functional genomics through high-quality reference genome assemblies.

The transition from non-threshable (hulled) to free-threshing grain was a defining event in wheat domestication, enabling the large-scale harvest and processing that underpinned early agricultural civilizations. Domesticated wheat can be classified either as hulled (DEW) or free-threshing (durum and bread wheat, plus other marginally cultivated primitive wheats). In hulled wheats, the grain is tightly enclosed by tough glumes that adhere firmly to the rachis, requiring additional dehulling steps prior to processing. Free-threshing wheats, by contrast, shed their grain readily upon threshing. Threshability is influenced by three main loci in wheat: the *Q* locus located on the long arm of chromosome 5A (Haas et al., 2019; Sharma et al., 2019), and two *Tenacious glume 1* (*Tg1*) loci placed on the short arms of chromosomes 2A and 2B. The fine mapping of the *Tg* loci revealed that dominant alleles of emmer wheat (*Tg1-A* and *Tg1-B*) determine, along with the recessive *Q* wild-type allele (qq), responsible for the non-free-threshing (tenacious glumes) phenotype (Faris et al., 2014; Simons et al., 2006; Sharma et al., 2019). Despite progress in characterizing these major determinants, a comprehensive molecular characterization of the principal threshability locus in tetraploid wheat, including its genomic architecture, expression dynamics, and functional basis, remains incomplete.

Crosses between free-threshing durum cultivars and WEW accessions generate segregating populations that are particularly informative for dissecting threshability genetics, as they capture the full range of variation at the *Q* and *Tg1* loci in a defined tetraploid background. Quantitative trait locus (QTL) analysis in such populations has identified genomic regions of large effect on glume tenacity and threshability, but the causal genes and molecular mechanisms underlying these QTLs have not been fully resolved. Near-isogenic lines (NILs) carrying small introgressions from WEW in an otherwise elite background provide a powerful complementary tool for validating QTL effects and isolating the functional variants responsible for phenotypic differences (Avni et al., 2018).

The objective of the current study is to identify genetic factors controlling free-threshing in domesticated wheat. To accomplish this, we conducted field experiments with a durum wheat × wild emmer recombinant inbred line (RIL) population across two environmental conditions, performed QTL analysis, and validated a major QTL controlling glume tenacity using a near-isogenic line (NIL) carrying a small introgression from WEW in the background of durum wheat Svevo. Our results provide new insight into the genetic architecture of threshability in tetraploid wheat and lay the groundwork for functional gene validation and marker-assisted breeding applications.

## 2. Materials and Methods

### 2.1. Tetraploid Wheat Mapping Population

A segregating population of 137 F7 recombinant inbred lines (RILs) developed via single-seed descent from a cross between the elite durum wheat cultivar ‘Svevo’ (Sv) and the wild emmer wheat (WEW) accession ‘Zavitan’ (Zv) was used for QTL mapping. The RIL population was advanced to the F7 generation to achieve near-complete homozygosity across the genome. All analyses were performed in conjunction with a previously developed high-density genetic map of this population (Avni et al., 2014).

### 2.2. Growing Conditions and Experimental Design

The Sv × Zv RIL population was phenotyped for threshability-related traits under field conditions in two independent environments in Israel. One experiment was conducted in 2014 at the Agricultural Research Organization experimental station in Rehovot (hereafter 2014R), and the second in 2015 at the coastal research station in Atlit (hereafter 2015A). Both experiments were arranged in a randomized complete block design (RCBD) with five replications; each experimental unit consisted of ten plants per genotype per replication. All 137 RILs and the two parental lines (Svevo and Zavitan) were included in both environments.

Supplemental irrigation was applied weekly when natural precipitation was insufficient. A slow-release fertilizer was applied at a rate of 50 kg ha^−1^ at sowing in both experiments. The two sites differ markedly in soil composition: the Atlit soil is a brown clay loam (organic matter, 1.7%; sand, 22.6%; silt, 57.0%; clay, 18.7%), whereas the Rehovot soil is a brown-red degraded sandy loam (Rhodoxeralf) composed of 76% sand, 8% silt, and 16% clay.

### 2.3. Phenotypic Evaluation of Threshability Traits

At maturity, three to six spikes were randomly selected from each experimental unit (genotype × replication combination) and used for phenotypic characterization. Three domestication-related traits were recorded (Table 1): glume tenacity (TG), assessed on a 1–5 scale by manual evaluation of glume adherence during threshing; seed number per spike (SDNPS), determined by counting all seeds from a representative spike; and the threshability ratio (THRR), calculated as the proportion of seeds that separated freely upon threshing (THR, Fig. 1, suplammentry video 1) relative to the total seed number per spike (THRR =THR / SDNPS).

**Table 1.** Domestication-related traits measured in the present study.

| Trait (Abbreviation) | Scale / Units | Measurement |
| --- | --- | --- |
| Glume tenacity (TG) | 1–5 scale | Glume texture evaluated by hand assessment of resistance during threshing |
| Seed number per spike (SDNPS) | Count (integer) | Total seed number counted from one representative spike |
| Threshability ratio (THRR) | Proportion (0–1) | Number of threshed seeds divided by total seed number per spike (THR / SDNPS) |

**Fig. 1.**
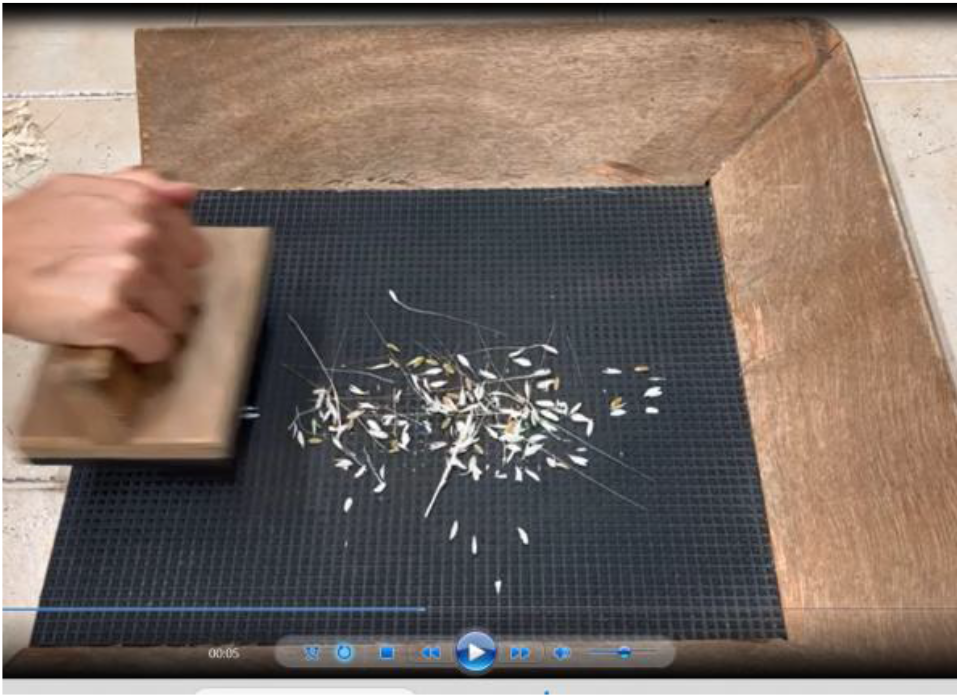
Measuring threshability in wheat using a threshing apparatus and parental line Svevo. Image from a video available at Supplementary Video 1. Trait values for each experimental unit were averaged across the three to six scored spikes prior to statistical analysis.

To complement the threshability data, we measured the glume pull-off force of the wheat genotypes using a mechanical force gauge (Wagner Force Dial FDK 4. Fig. 2).

**Fig. 2.**
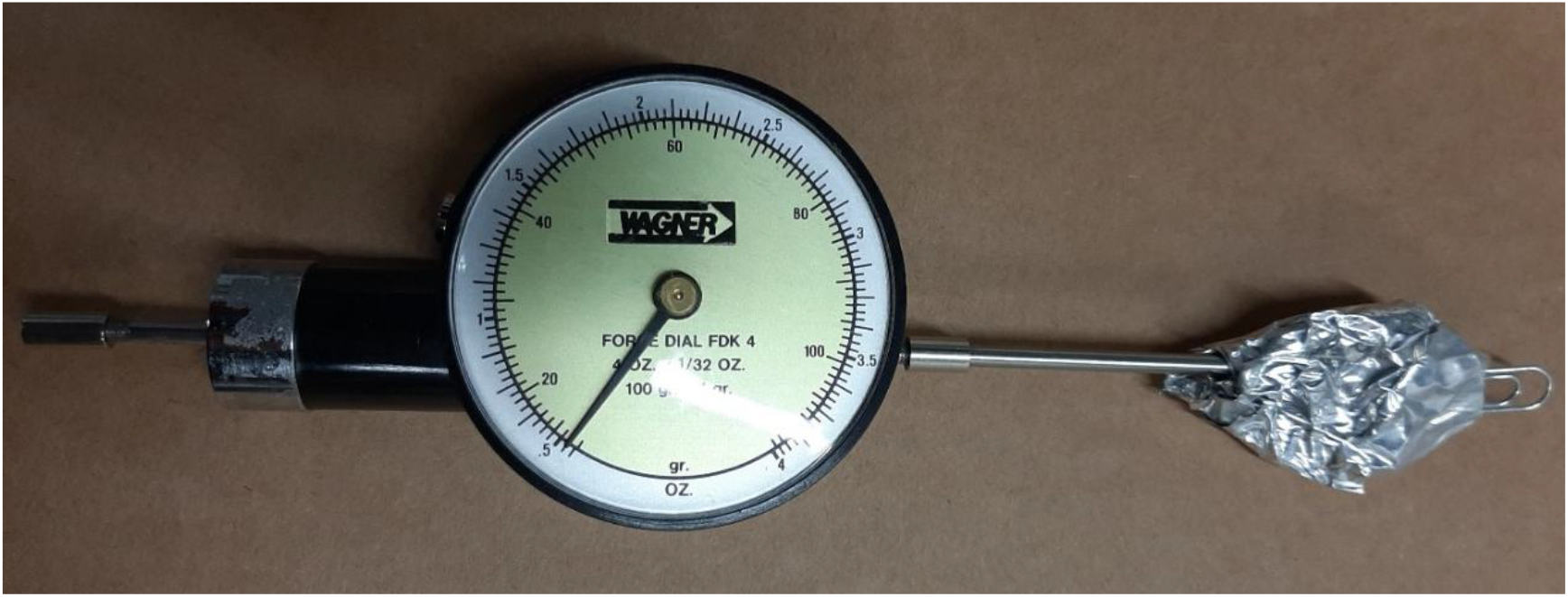
Measurement of glume pull-off force. A mechanical force gauge (Wagner Force Dial FDK 4) configured to quantify the detachment force required to remove the glume from the rachis.

### 2.4. QTL Analysis

QTL analysis was performed using the MultiQTL software package following the procedures described in Nave et al., 2016. Interval mapping was conducted using the high-density Sv × Zv genetic map, and the significance threshold for each QTL was determined by a permutation test (1,000 permutations; α = 0.05). Detected QTLs were subsequently subjected to genotype × environment (G × E) interaction analysis by ANOVA to assess the consistency of QTL effects across the two growing environments.

### 2.5. Development and Evaluation of Near Isogenic Line NIL-65

To validate the major threshability QTL identified in the RIL population, RIL-65, which carried the WEW allele at the target locus (*Tg1-B*), was selected as a donor and backcrossed three times to the recurrent parent Svevo (BC3). Following the third backcross, selected plants were self-pollinated for five consecutive generations to achieve homozygosity. A single BC_3_F_5_ Near Isogenic Line (NIL-65) was identified and confirmed to carry a homozygous WEW introgression at the target region by low-coverage whole-genome skim sequencing, enabling precise delineation of introgression boundaries in the Svevo genetic background.

## 3. Results

### 3.1. Phenotypic Variation in the Sv × Zv RIL Population

Phenotypic distributions for all threshability-related traits scored across two environments (2014 and 2015) are presented in Fig. 3. The parental lines differed markedly for all three traits. Svevo displayed high seed number per spike (SDNPS ∼75 in 2014, ∼42 in 2015), high threshability (THRR ∼0.96 and ∼0.98, respectively), and low glume tenacity (TG score 1 in both years), consistent with its free-threshing durum phenotype. Zavitan, by contrast, exhibited lower seed number per spike (SDNPS ∼35 and ∼31), near-zero threshability (THRR ∼0.01 and ∼0.02), and maximum glume tenacity (TG score 5 in both years), consistent with its hulled, non-free-threshing wild emmer phenotype (Table S1).

**Fig. 3.**
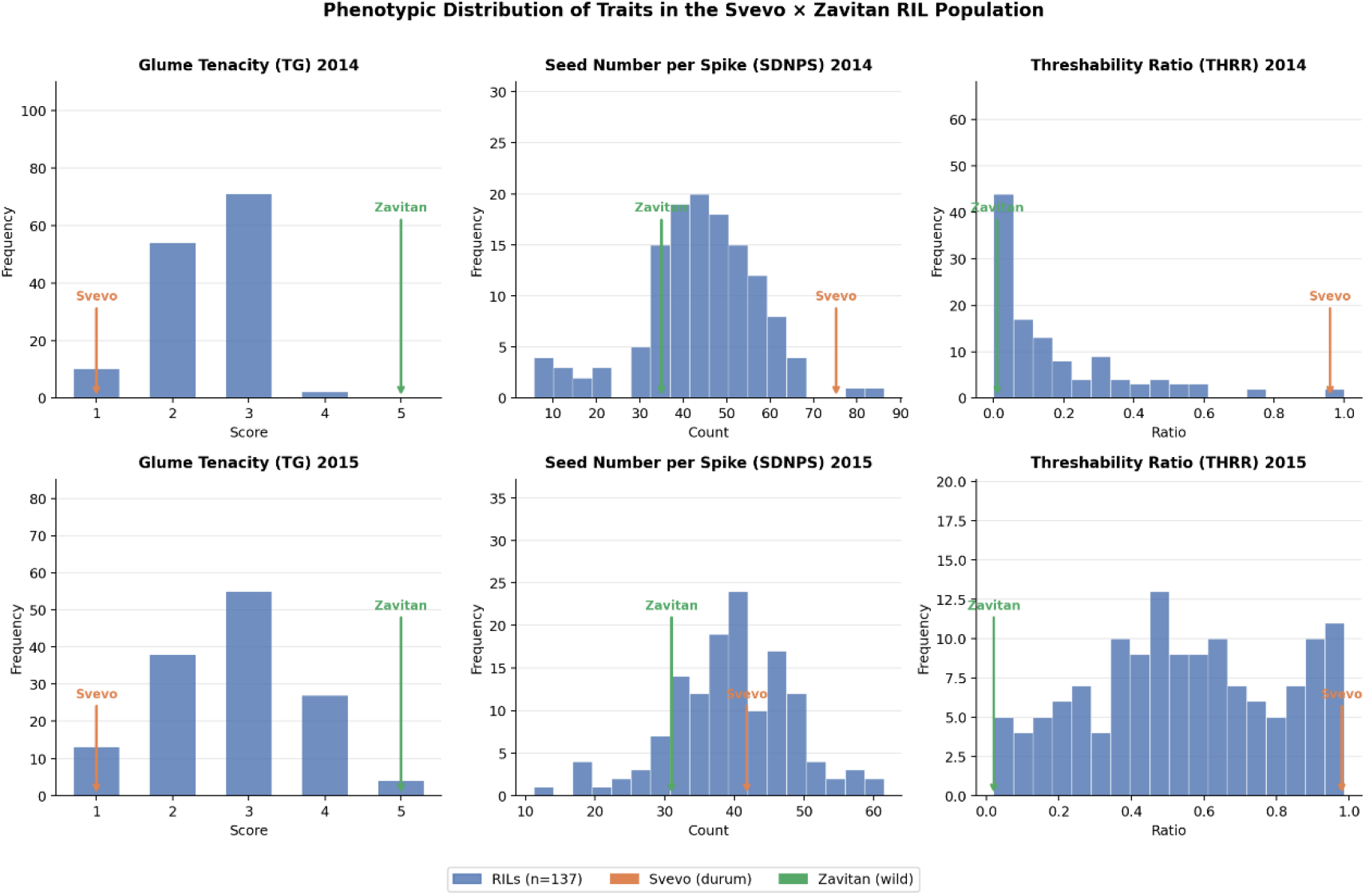
Phenotypic distributions of threshability-related traits in the Svevo × Zavitan recombinant inbred line (RIL) population. Frequency distributions of glume tenacity (TG), seed number per spike (SDNPS), and threshability ratio (THRR) measured across 137 RILs in 2014 (top row) and 2015 (bottom row). TG was scored on a discrete 1–5 scale (1 = free-threshing, 5 = fully tenacious). THRR represents the proportion of seeds released by mechanical threshing relative to total seed number, and SDNPS reflects total seed set per spike. Colored arrows indicate the phenotypic values of the parental lines: Svevo (durum wheat, orange) and Zavitan (wild emmer wheat, green). The contrasting parental values for TG and THRR reflect the divergent threshability phenotypes that drive segregation in this population.

Among the RILs, SDNPS followed an approximately normal distribution in both years, centered around 35–50 seeds per spike, with transgressive segregants extending both below the Zavitan mean and toward the Svevo range. THRR displayed a predominantly bimodal distribution: the majority of RILs clustered near zero, resembling the hulled Zavitan parent, while a smaller class segregated toward a free-threshing phenotype (THRR ∼1.0), suggesting that threshability in this population tends toward a discrete, threshold-like trait rather than a strictly continuous one. Glume tenacity (TG) showed a similarly skewed distribution, with most RILs scoring in the lower range (TG 1–2), while a subset exhibited higher tenacity scores (TG 3–5) comparable to Zavitan. The non-uniform distributions of THRR and TG are consistent with the possibility that one or a few loci of relatively large effect contribute substantially to the segregation of threshability in the Sv × Zv population, a hypothesis explored further in the QTL analysis below.

### 3.2. QTL Analysis

QTL analysis across the two environments (2014R and 2015A) identified a total of 19 significant QTLs distributed across six chromosomes for the three traits examined (Table 2). Where corresponding QTLs were detected in both environments, these are treated as a single environmentally stable locus. The genomic positions, LOD scores, proportion of explained variance (PEV), and allelic effects of all detected QTLs are summarized in Table 2.

**Table 2.** QTLs detected for threshability-related traits in the Svevo × Zavitan recombinant inbred line population (n = 137) across two field environments in Israel (2014R and 2015A). Traits scored were seed number per spike (SDNPS), glume tenacity (TG), and threshability ratio (THRR). For each QTL, the chromosome (Chr), environment, LOD score, peak position (cM), proportion of explained variance (PEV, %), estimated additive effect, and favorable allele are indicated. QTLs detected in both environments at the same chromosomal region are considered environmentally stable loci.

| Trait | Chr | Year | LOD<sup>1</sup> | Position (cM) | PEV<sup>2</sup> (%) | Estimated effect<sup>3</sup> | Favourable Allele<sup>4</sup> |
| --- | --- | --- | --- | --- | --- | --- | --- |
| SDNPS | 1B | 2015A | 2.47 | 97.59±25.39 | 0.053±0.024 | 3.405±2.037 | S |
|  | 4B | 2015A | 1.93 | 65.13±21.38 | 0.052±0.024 | -2.817±2.804 | Z |
|  | 5A | 2014R | 5 | 43.09±12.89 | 0.157±0.051 | 10.36±4.915 | S |
|  | 5A | 2015A | 6.76 | 41.99±7.968 | 0.196±0.052 | 7.624±1.448 | S |
|  | 6A | 2015A | 3.87 | 69.09±11.24 | 0.087±0.031 | 4.961±1.269 | S |
|  | 7A | 2015A | 2.64 | 105.28±32.62 | 0.069±0.029 | 3.941±2.317 | S |
| TG | 2A | 2014R | 5.06 | 29.7±17.62 | 0.11±0.032 | -0.42±0.071 | Z |
|  | 2A | 2015A | 3.62 | 27.03±14.47 | 0.097±0.029 | -0.59±0.115 | Z |
|  | 2B | 2014R | 6.02 | 42.39±7.899 | 0.134±0.041 | -0.464±0.088 | Z |
|  | 2B | 2015A | 8.98 | 45.47±6.79 | 0.198±0.044 | -0.844±0.115 | Z |
|  | 4B | 2014R | 4.55 | 52.85±23.29 | 0.105±0.035 | -0.308±0.274 | Z |
|  | 4B | 2015A | 4.97 | 48.09±13.73 | 0.1±0.032 | -0.548±0.254 | Z |
|  | 5A | 2014R | 6.57 | 173.17±9.753 | 0.15±0.042 | -0.485±0.089 | Z |
|  | 5A | 2015A | 5.56 | 168.88±10.89 | 0.124±0.038 | -0.666±0.123 | Z |
| THRR | 2A | 2014R | 3.2 | 54.01±30.1 | 0.129±0.034 | 0.152±0.024 | S |
|  | 2A | 2015A | 8.2 | 31.8±7.732 | 0.196±0.046 | 0.225±0.033 | S |
|  | 2B | 2014R | 6.43 | 44.61±16.43 | 0.227±0.07 | 0.194±0.057 | S |
|  | 2B | 2015A | 14.22 | 44.4±1.935 | 0.312±0.046 | 0.285±0.029 | S |
|  | 4B | 2015A | 3.73 | 44.89±18.51 | 0.06±0.023 | 0.101±0.074 | S |

<sup>1</sup>LOD scores that were found significant when comparing H0:H1 (there is significant QTL/s on the chromosome Vs no significant effect of the chromosome) or H1:H2 in case of significant two-linked QTLs using 10,000 permutation tests applied after MIM analysis.
<sup>2</sup>Proportion of the explained variance of the trait
<sup>3</sup>The adaptive effect of an allele calculated as one half of the mean differences between homozygote genotypes with the allele and without it. (+ the trait avg is higher with the Sv allele, - the avg is higher with the Zv allele).
<sup>4</sup>The parental allele that contributes to higher values of the measured trait.

#### Seed Number Per Spike (SDNPS)

Six significant QTLs were detected for SDNPS, with LOD scores ranging from 1.93 to 6.76 and individually explaining 5.2–19.6% of the phenotypic variance. The most prominent QTL mapped to chromosome 5A and was detected in both environments (LOD 5.00–6.76; PEV 15.7–19.6%), making it the most consistent and largest-effect locus for this trait. At five of the six loci (1B, 5A, 6A, and 7A), the Svevo allele conferred higher seed number. The exception was the 4B QTL, where the Zavitan allele was associated with higher SDNPS values, suggesting that the WEW parent carries favorable alleles at this locus despite its overall lower seed number phenotype.

#### Glume Tenacity (TG)

Eight significant QTLs were detected for TG, with LOD scores ranging from 3.62 to 8.98 and explaining 9.7–19.8% of the phenotypic variance per locus. QTLs mapped to chromosomes 2A, 2B, 4B, and 5A, and were consistently detected in both environments at each locus, indicating robust and environmentally stable genetic effects. At all eight loci, the Zavitan allele conferred higher glume tenacity, consistent with the hulled, non-free-threshing phenotype of WEW. Notably, the 5A QTL peak co-localized with the *Q* locus marker (*Q-5A*), implicating this well-characterized domestication gene in the segregation of TG in this population. The strongest single-environment effect was observed on chromosome 2B in 2015A (LOD 8.98; PEV 19.8%).

#### Threshability Ratio (THRR)

Five significant QTLs were detected for THRR, with LOD scores ranging from 3.20 to 14.22 and explaining 6.0–31.2% of the phenotypic variance. QTLs mapped to chromosomes 2A, 2B, and 4B. At all five loci, the Svevo allele conferred higher threshability, consistent with its free-threshing phenotype. The largest-effect QTL mapped to chromosome 2B and was detected in both environments (LOD 6.43–14.22; PEV 22.7–31.2%). Because THRR is ratio-based and SDNPS varies genetically, we interpret this peak alongside co-localized TG signals and mechanical pull-off force differences, which together indicate that the 2B region is the principal determinant of threshability in this cross. As expected, the chromosomal positions of the THRR QTLs on 2A and 2B co-localized with the TG QTLs on the same chromosomes, suggesting that glume tenacity and threshability are controlled by the same, or tightly linked, genomic regions, consistent with their biological relationship as components of the same threshing phenotype.

### 3.3. Development and Validation of Near-Isogenic Line NIL-65

The QTL analysis identified a major locus for glume tenacity (TG) on the short arm of chromosome 2B, detected in both environments (LOD 6.02–8.98; PEV 13.4–19.8%), with the Zavitan allele conferring higher glume tenacity at this locus. Physical mapping using flanking markers IWB2315 and IWB21536 (Fig. 4) delimited the QTL to a ∼5.8 Mbp interval in the Zavitan Rel. 2 reference assembly (Zhu et al., 2019) (Chr. 2B: 37,213,853–46,678,663), corresponding to the previously described *Tg1-B* locus.

**Fig. 4.**
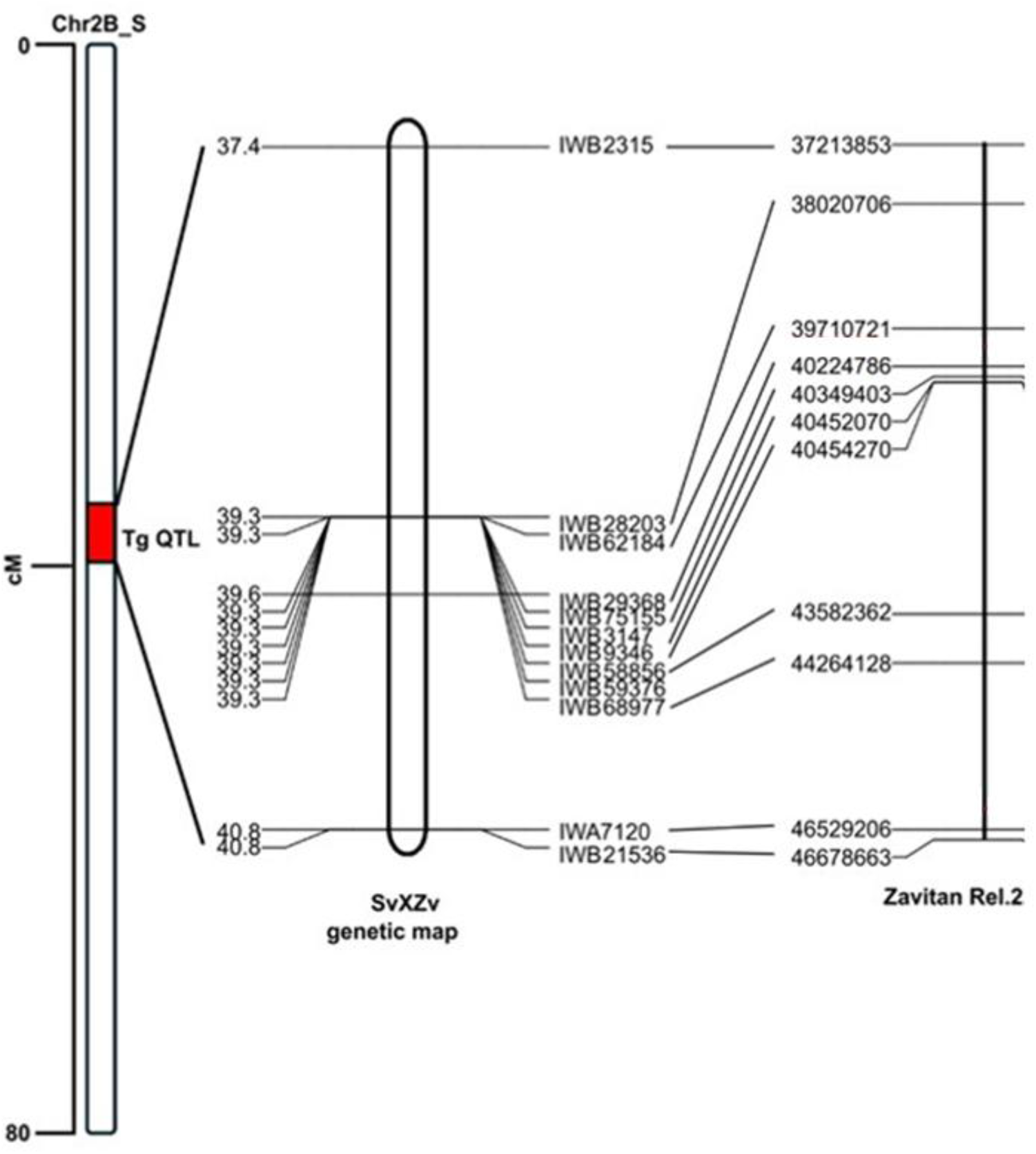
Genetic and physical mapping of the *Tg1-B* locus on chromosome 2B. Schematic representation of the *Tg1-B* QTL interval on the short arm of chromosome 2B (Chr2B_S), shown alongside the corresponding region of the Svevo × Zavitan (SvXZv) genetic map. Genetic markers are connected to their physical positions on the Zavitan Rel. 2 genome assembly (right, in bp). The *Tg1-B* QTL (red bar) maps to a ∼3.4 cM interval between 37.4 and 40.8 cM. Flanking markers IWB2315 define the proximal boundary of the interval, while IWA7120 and IWB21536 define the distal boundary.

In the Svevo Rel.1.0 reference assembly (Maccaferri et al., 2019), some of the proximal markers localized to unanchored contigs on the unmapped chromosome (chrUn), precluding precise determination of the QTL interval in the durum background.

To validate the effect of the chromosome 2B *Tg1-B* QTL independently of the broader RIL genetic background, RIL-65, which carries the Zavitan allele at this locus and exhibits the tenacious glume phenotype, was selected as a donor and backcrossed three times to the recurrent parent Svevo, which carries the susceptible *tg1-B* allele. Progeny were selected at each backcross generation for retention of the target introgressions by using markers IWB2315 and IWB21536 that flank the *Tg1-B* QTL. The resulting BC_3_ plants were self-pollinated for five generations to produce the near-isogenic line NIL-65 (BC_3_F_5_). Low-coverage whole-genome skim sequencing of NIL-65 confirmed the presence of a homozygous Zavitan introgression encompassing the *Tg1-B* locus on chromosome 2B (Fig. 5). In addition to the target introgression, skim sequencing revealed seven residual WEW introgressions distributed across the genome (Fig. 5). Importantly, none of these residual introgressions overlapped with any previously mapped threshability QTL, indicating that NIL-65 can be considered a reliable stocks line for the *Tg1-B* WEW allele.

**Fig. 5.**
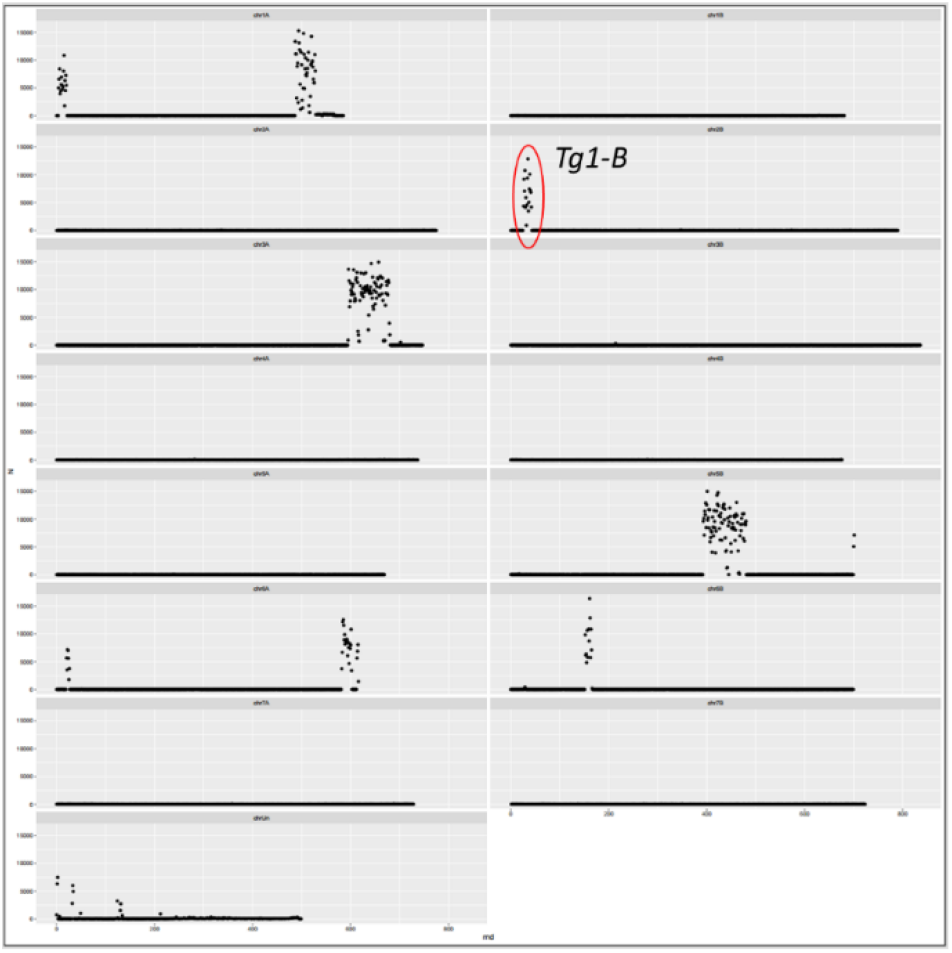
Whole-genome skim sequencing profile of NIL-65. Chromosomal distribution of Zavitan introgressions against the Svevo genetic background. The target *Tg1-B* introgression on chromosome 2B (indicated in red), along with seven additional residual WEW introgressions retained following backcrossing.

To further reduce residual background introgressions, a segregating BC_4_F_2_ population was developed by crossing NIL-65 to Svevo. Glume pulling force was measured across this population using a mechanical force gauge (Fig. 6, Table S2), revealing a continuous phenotypic distribution consistent with segregation at a major locus in an F_2_ population.

**Fig. 6.**
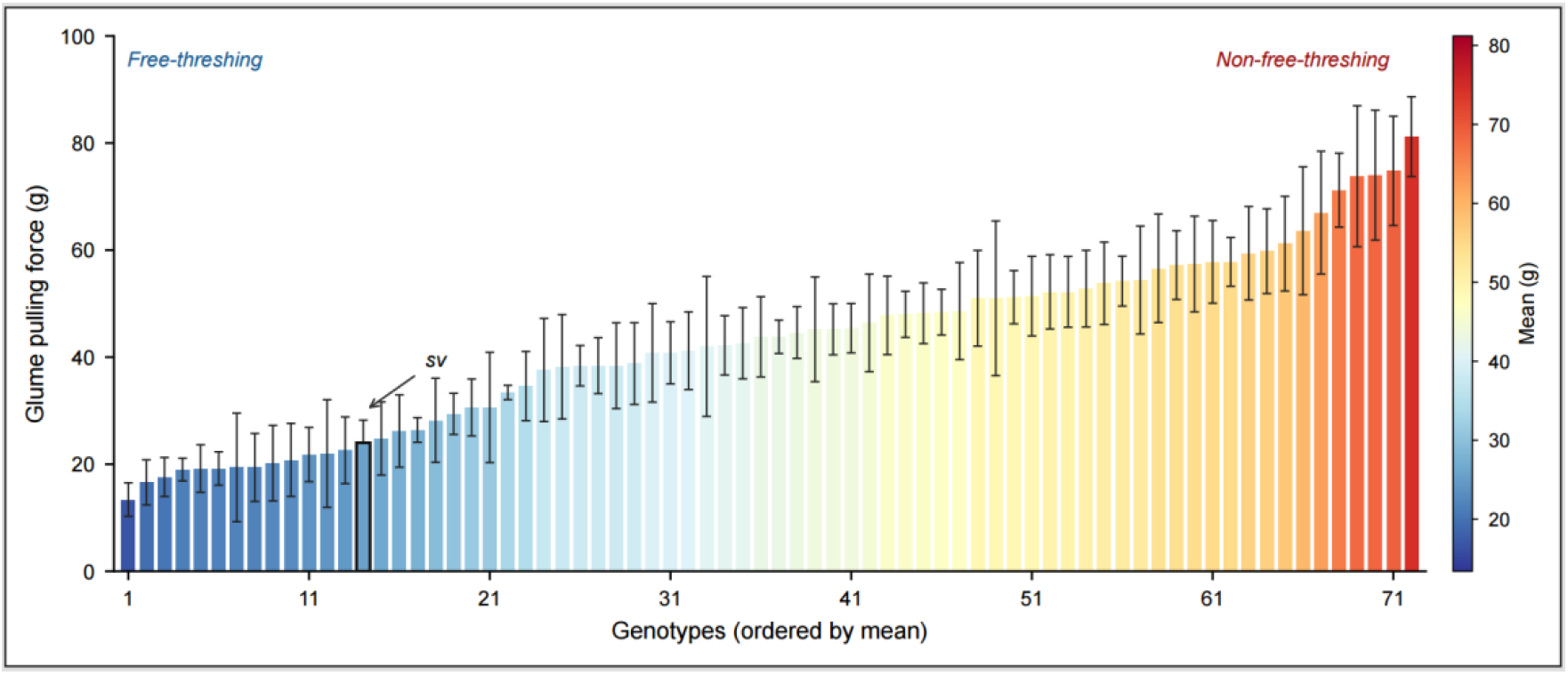
Phenotypic distribution of glume pulling force (g) in the NIL-65 × Svevo BC_4_F_2_ population and parental line Svevo (indicated by arrow). Each data point represents the mean of five replicate measurements per plant.

To investigate genotype-phenotype relationships, BC_4_F_2_ individuals with extreme threshability phenotypes were selected for genotyping-by-sequencing (GBS)-based Bulk Segregant Analysis (BSA). Two contrasting bulks were established: 13 plants with the highest glume pulling force (mean 65.9 g; Table S3), representing the tenacious-glume class, and 12 plants with the lowest glume pulling force (mean 19.3 g; Table S3), representing the free-threshing class. The two bulks differed highly significantly in glume tenacity (Welch’s t-test, p < 0.001). Sequence read-depth analysis revealed a ∼11 Mb Zavitan-derived introgression on chromosome 2B, spanning the *Tg1-B* locus, that was present exclusively in the tenacious-glume bulk and absent from the free-threshing bulk (Fig. 6; Table S4), pinpointing this interval as the primary determinant of threshability segregation in the BC_4_F_2_ population.

None of the seven residual introgressions showed a statistically significant association with glume pulling force in the BC_4_F_2_ population (p > 0.01 for all; Supplementary Table S4), whereas the chromosome 2B *Tg1-B* introgression remained the sole genomic region significantly associated with the tenacious-glume phenotype. Notably, one introgression initially mapped to an unanchored chromosome (Chr U) using the Svevo Rel.1.0 (Maccaferri et al., 2019) was subsequently localized to chromosome 2B upon remapping against both the Zavitan Rel. 2 genome reference (Zhu et al., 2019) and the Svevo Rel.2.0 reference (Mazzucotelliet al. submitted), confirming that it represents an additional segment of the same chromosomal region rather than an independent locus. These results demonstrate that the phenotypic contrast between *Tg1-B* and *tg1-B* sister lines is attributable specifically to the chromosome 2B introgression and cannot be explained by co-segregation of any other residual WEW segment.

Consistent with this, visual inspection revealed that plants carrying *Tg1-B* maintained structurally intact glumes, whereas *tg1-B* sister lines exhibited characteristic cracks at the glume base, a morphological feature associated with the free-threshing phenotype (Fig. 7).

**Fig. 7.**
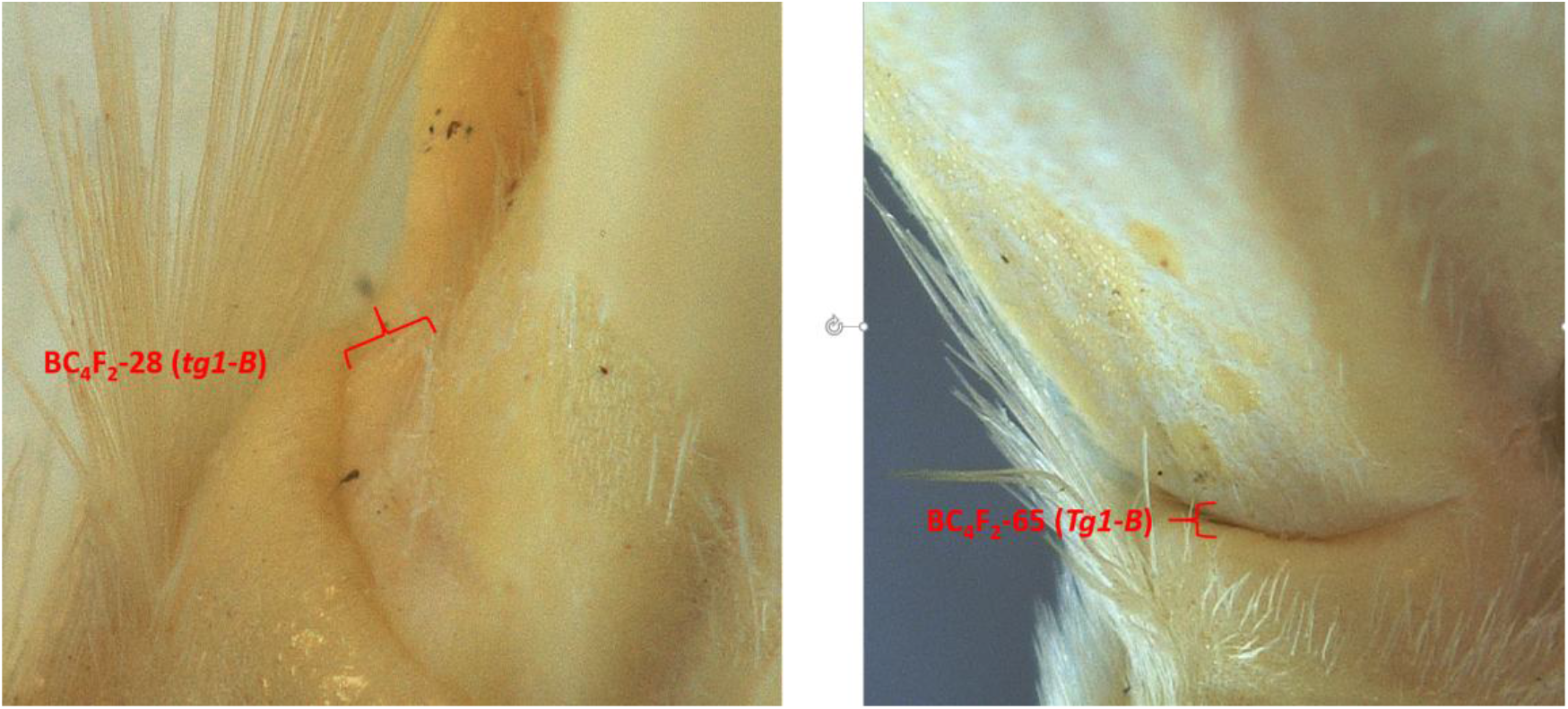
Microscopic images of glume bases in the NIL-65 × Svevo BC_4_F_2_ population. Plants carrying *Tg1-B* (e.g., BC_4_F_2_-65; right) display structurally intact, firmly attached glumes, whereas *tg1-B* sister lines (e.g., BC_4_F_2_-28; left) exhibit characteristic fractures at the glume base. This structural breakdown, particularly at the glume attachment zone, likely reflects targeted tissue weakening that reduces the mechanical force required for grain release; a defining morphological hallmark of the free-threshing phenotype.

## 4. Discussion

### 4.1. The *Tg1-B* Locus Is the Principal Determinant of Glume Tenacity and Threshability in the Svevo × Zavitan Population

QTL analysis across two field environments revealed a highly consistent and large-effect locus for both glume tenacity and threshability on chromosome 2B, corresponding to the *Tg1-B* locus previously identified in tetraploid wheat (Faris et al., 2014; Sharma et al., 2019). The co-localization of TG and THRR QTLs at the same chromosomal position on 2B, and again on 2A (Table 2), is consistent with the view that glume tenacity and threshability are physiologically coupled traits: the mechanical strength of the glume-rachis junction directly determines the ease with which grain is released during threshing. The detection of a 5A QTL that co-localizes with the *Q* marker in the TG analysis provides further evidence that this well-characterized domestication gene contributes to the glume tenacity phenotype in a tetraploid background, consistent with its described role as a pleiotropic regulator of spike morphology and threshability (Haas et al., 2019; Sharma et al., 2019).

### 4.2. NIL-65 Validates the *Tg1-B* QTL and Provides a Defined Genetic Resource for Gene Discovery

The development and phenotypic evaluation of NIL-65 confirms that the *Tg1-B* QTL effect is reproducible independently of the broader RIL genetic background. The clear segregation of glume pulling force in the BC_4_F_2_ population, and the significant difference in threshability between plants homozygous for the *Tg1-B* versus *tg1-B* alleles, demonstrate that a single introgression from WEW is sufficient to convert a free-threshing durum background to a tenacious-glume phenotype (Fig. 6). This is consistent with the dominant inheritance model described for *Tg1* loci by Faris et al. (2014), and reinforces the interpretation that the WEW-derived allele acts as a gain-of-function determinant of glume tenacity.

The seven residual WEW introgressions detected in NIL-65 by skim sequencing (Fig. 5) represent a limitation inherent to NIL development from wide crosses. However, none of these introgressions showed a statistically significant association with glume pulling force in the BC_4_F_2_ population (p > 0.01 for all; Supplementary Table S4), nor did any overlap with previously mapped threshability QTL. Collectively, these findings support the use of NIL-65 as a reliable genetic stock for functional characterization of the *Tg1-B* WEW allele.

### 4.3. Morphological Basis of Glume Tenacity: Glume Cracking as a Marker of the Free-Threshing State

The visual phenotyping data provide important insight into the cellular and structural basis of glume tenacity (Fig. 7). The characteristic basal cracking observed in *tg1-B* lines suggests that the free-threshing phenotype arises, at least in part, from structural weakening of the abscission zone at the base of the glume. This is consistent with previous histological and transcriptomic studies indicating that cell wall modification and reduced lignification at the glume-rachis junction facilitate grain separation in free-threshing wheats (Debernardi et al., 2017). By contrast, the intact glume morphology of *Tg1-B* plants implies that the WEW allele maintains the structural integrity of this zone, either by reinforcing cell wall composition or by suppressing the abscission-related developmental program. Future histological analysis of glume base tissue in NIL-65 and its *tg1-B* sister lines would be valuable for testing this hypothesis and for identifying the cell types and developmental stages at which *Tg1-B* exerts its effect.

### 4.4. Implications for Candidate Gene Identification and Breeding Applications

The approximately 11 Mb introgression delimited by GBS in the BC_4_F_2_ population provides a refined physical interval for candidate gene prioritization. The availability of the high-quality Zavitan WEWSeq v2.0 and Svevo Rel.2.0 reference assemblies now enables systematic comparison of the gene content and allelic variants within this interval between the hulled and free-threshing haplotypes. Priority candidates include genes involved in cell wall biosynthesis and remodeling, transcription factors expressed in the developing spike, and any gene showing differential expression between NIL-65 and its *tg1-B* sister lines.

From an applied perspective, the NIL-65 germplasm and the flanking markers defined in this study provide immediate tools for marker-assisted selection (MAS) targeting glume tenacity. While the free-threshing trait is agronomically desirable in modern durum and bread wheat breeding, precise manipulation of threshability, including the ability to engineer intermediate phenotypes or to restore glume tenacity in specialty grain markets, requires detailed knowledge of the underlying genes and their regulatory networks. The near-isogenic materials generated here, alongside the refined physical interval on chromosome 2B, constitute a foundation for the next phase of functional genomic investigation into one of the most consequential traits of wheat domestication.

## Supporting information

Supplemental video 1 NIL-65

Supplemental video 1 Svevo

## Funding

This research was funded the Israel Science Foundation (ISF grants 1431/22 and 2223/22).

## Acknowledgment

We thank Dr. Tami Krugman for valuable discussions throughout this work, and take this opportunity to congratulate her on her forthcoming retirement after four decades of distinguished contributions to cereal research.

## Competing interests

The authors declare no competing interests.

## Notes

### Competing Interest Statement

The authors have declared no competing interest.

